# Epigenetic regulator miRNA pattern differences among SARS-CoV, SARS-CoV-2 and SARS-CoV-2 world-wide isolates delineated the mystery behind the epic pathogenicity and distinct clinical characteristics of pandemic COVID-19

**DOI:** 10.1101/2020.05.06.081026

**Authors:** Md. Abdullah-Al-Kamran Khan, Md. Rabi Us Sany, Md. Shafiqul Islam, Md. Saheb Mehebub, Abul Bashar Mir Md. Khademul Islam

**Affiliations:** Department of Mathematics and Natural Sciences, BRAC University, Dhaka, Bangladesh; Department of Genetic Engineering & Biotechnology, University of Dhaka, Dhaka, Bangladesh

**Author notes:** Correspondence: Dr. Abul Bashar Mir Md. Khademul Islam, Associate Professor, Department of Genetic Engineering and Biotechnology, University of Dhaka, Dhaka 1000, Bangladesh.

**Keywords:** SARS-CoV-2, COVID-19, miRNA-microRNA, viral pathogenesis, immune regulation

## Abstract

Detailed molecular mechanism of SARS-CoV-2 pathogenesis is still elusive to address its deadlier nature and to design effective theraputics. Here, we present our study elucidating the interplay between the SARS-CoV and SARS-CoV-2 viruses’; and host’s miRNAs, an epigenetic regulator, as a mode of pathogenesis, and enlightened how the SARS-CoV and SARS-CoV-2 infections differ in terms of their miRNA mediated interactions with host and its implications in the disease complexity. We have utilized computational approaches to predict potential host and viral miRNAs, and their possible roles in different important functional pathways. We have identified several putative host antiviral miRNAs that can target the SARS viruses, and also SARS viruses’ encoded miRNAs targeting host genes. *In silico* predicted targets were also integrated with SARS infected human cells microarray and RNA-seq gene expression data. Comparison of the host miRNA binding profiles on 67 different SARS-CoV-2 genomes from 24 different countries with respective country’s normalized death count surprisingly uncovered some miRNA clusters which are associated with increased death rates. We have found that induced cellular miRNAs can be both a boon and a bane to the host immunity, as they have possible roles in neutralizing the viral threat, parallelly, they can also function as proviral factors. On the other hand, from over representation analysis, interestingly our study revealed that although both SARS-CoV and SARS-CoV-2 viral miRNAs could target broad immune signaling pathways; only some of the SARS-CoV-2 miRNAs are found to uniquely target some immune signaling pathways like-autophagy, IFN-I signaling etc, which might suggest their immune-escape mechanisms for prolonged latency inside some hosts without any symptoms of COVID-19. Further, SARS-CoV-2 can modulate several important cellular pathways which might lead to the increased anomalies in patients with comorbidities like-cardiovascular diseases, diabetes, breathing complications, etc. This might suggest that miRNAs can be a key epigenetic modulator behind the overcomplications amongst the COVID-19 patients. Our results support that miRNAs of host and SARS-CoV-2 can indeed play a role in the pathogenesis which can be further concluded with more experiments. These results will also be useful in designing RNA therapeutics to alleviate the complications from COVID-19.

## 1. Introduction

Coronavirus outbreaks have been reported over the past three decades, but the recent SARS-CoV-2 pandemic has outreached more than 200 countries and has been the causative agent for the death of 58,392 people around the globe and 1,087,374 coronavirus cases have been filed till the date of writing this article (Worldometer, 2020). Among closed cases of SARS-CoV-2, 20% of the patients have died and 5% of patients within active cases are in critical situations (Worldometer, 2020). The initial estimation of SARS-CoV-2 death rate was 3.4%, declared by WHO (WHO, 2020) requires a refresh as the global casualty is uprising, thus, this novel virus requires novel and in-depth studies to promote new strategies for the management of this pandemic.

Coronavirus subfamily is a single-stranded positive-sense (+ssRNA) virus with a genome size of around 30kb (Lu et al., 2020). The family is categorized into four subgenera as alpha, beta, gamma, and delta coronavirus (Cheng and Shan, 2020). SARS-CoV-2 is a beta coronavirus with a genome size of 29.9kb (Accession no. NC_045512.2) with 11 genes being reported in NCBI Gene (NCBI-Gene, 2020). Phylogenetic analysis between SARS-CoV-2 and SARS-CoV showed ∼79% similarity. Whereas the distance is much longer for MERS-CoV (∼50% similarity) but the closest relative to the SARS-CoV-2 is bat derived SARS-like coronavirus (∼90% similarity) (Jiang et al., 2020; Lu et al., 2020; Ren et al., 2020). Genomic analysis of SARS-CoV and SARS-CoV-2 has shown substitution of 380 amino acids and deletion of ORF8a, elongation of ORF8b (84 vs 121 amino acid residues) and truncation of ORF3b (154aa in SARS-CoV whereas 22aa in SARS-CoV) (Lu et al., 2020).

MicroRNAs are small ncRNA molecules that regulate post-transcriptional level gene expression and its already established that viruses use host machinery to produce miRNAs (Ambros, 2001). Although miRNA can be an important anti-viral tool (Trobaugh and Klimstra, 2017) which can stimulate the innate and adaptive immune system, (Ambros, 2001; Trobaugh and Klimstra, 2017) but also can be a back door for viral propagation due to being non-antigenic thereby modulating cellular pathways without triggering host immune response, (Cullen, 2013; Globinska et al., 2014) for example, nucleocapsid protein of coronavirus OC43 binds miR-9 and activates NF-κB (Lai et al., 2014). Although host microRNAs are either utilized or regulated by viruses, viral miRNAs are another side of the coin, where they regulate host gene expression, cellular proliferation, stress-related genes and even viral gene expression (Cullen, 2010; Haasnoot and Berkhout, 2011; Lai et al., 2014). A summary discussed that number of DNA and RNA viruses produce miRNAs known as viral miRNAs (v-miRNAs) to evade the host immune response (Mishra et al., 2020). Novel viral miRNAs have been predicted to play an important role in neurological disorders as well (Islam et al., 2019). Among RNA viruses, for example, HIV-1 encoded miR-H1 can cause mononuclear cells apoptosis; H5N1 influenza virus-encoded miR-HA-3p targets host PCBP2 and contributes to ‘cytokine storm’ and mortality, and KUN-miR-1 of West Nile virus targets host’s GATA4 which facilitates virus replication (Li and Zou, 2019). Host miRNAs interaction with SARS-CoV genome and viral proteins have been elucidated to suppress viral growth and immune evasion (Mallick et al., 2009). Novel classes of ncRNAs have been also observed by studies those might play a definitive role in pathogenesis and survival (Liu et al., 2018). Respiratory viral infections caused by influenza, rhinovirus, adenovirus, RSV and coronaviruses can be related to aberrant host miRNA expression and their effect on host can be like - cell apoptosis, inhibition of immunologic pathways, down regulation of host antiviral responses etc (Mallick et al., 2009; Bondanese et al., 2014; Islam et al., 2019; Li and Zou, 2019; Mishra et al., 2020). Transmissible gastroenteritis virus (TGEV) although induce significant IFN-I production after infection by inducing endoplasmic reticulum (ER), it can evade antiviral effect of IFN-I by downregulating miR-30a-5p that normally enhances IFN-I antiviral activity (Ma et al., 2018).

On the other hand, host miRNA expression plays a major role in controlling viral pathogenesis by mediating T cells and antiviral effector functions (Dickey et al., 2016). The first reported example of a cellular miRNA that targets a viral RNA genome is miR-32 which targets the retrovirus PFV-1 transcripts and results in reduced virus replication (Lecellier et al., 2005). Similarly, miR-24, miR-93 can target VSV virus L and P protein (Otsuka et al., 2007); miR-29a targets HIV Nef protein (Ahluwalia et al., 2008) to inhibit replication; miR-1, miR-30, miR-128, miR-196, miR-296, miR-351, miR-431, miR-448 targets HCV C and NS5A protein to inhibits translation/replication by inducing IFN signaling (Pedersen et al., 2007). Thus, miRNA can provide a different perspective in explaining the pathogenesis and infectivity of the novel SARS-CoV-2. Although SARS-CoV is distantly related to SARS-CoV-2, there are some similarities in their signs and symptoms even they might be similar in pathogenesis but there are crucial differences between two diseases too (Xu et al., 2020). On the other hand, SARS-CoV-2 has infected many countries, and which resulted a stable mutation rate and resulted some variation (Cullen, 2006; Dykxhoorn, 2007). There are evidence that viral pathogens can have novel immune evasion role by utilizing host miRNA (Islam et al., 2019; Mishra et al., 2020).

However, detailed miRNA mediated epigenetic interplay between SARS-CoV-2 and host yet to be elucidated. It is not known what the probable miRNAs produced by SARS-CoV-2 are affecting which human processes; also, which anti-viral miRNAs taking part in host immunity. The genomic difference which in result controls the host miRNA target sites and viral miRNAs might explain the difference SARS-CoV and various isolated of SARS-CoV-2 in pathogenesis and infectivity. Here in this study, we hypothesize on three potential effects of host and viral miRNA – (1) Genomic differences between SARS-CoV and SARS-CoV2 can led to variations in host miRNA binding and differences in hence pathogenicity, signs and symptoms of these diseases and might explain the relatively longer incubation period of SARS-CoV-2. (2) Similarly, on the other hand, there might be differences in viral miRNAs that can regulate expressions of different sets of host genes which in turn can be advantageous to the virus or the host. (3) Due to fast mutation rate, observed variations among SARS-CoV-2 isolates in different regions of the world might result in variation in host capacities to target the virus with its miRNAs. This, in turn, might play a significant role in varying degrees of disease severity, symptoms and mortality rate in different regions. In this study, we have done comparative analysis between SARS-CoV and SARS-CoV-2 in respect of host miRNA–viral genome interaction as well their differences based on region-specific isolates of SARS-CoV-2; viral miRNA-host mRNA interactions to delineate the exclusive features of COVID-19 and their roles in viral survival and pathogenicity in respect of SARS-CoV (Figure 1).

**Figure 1:**
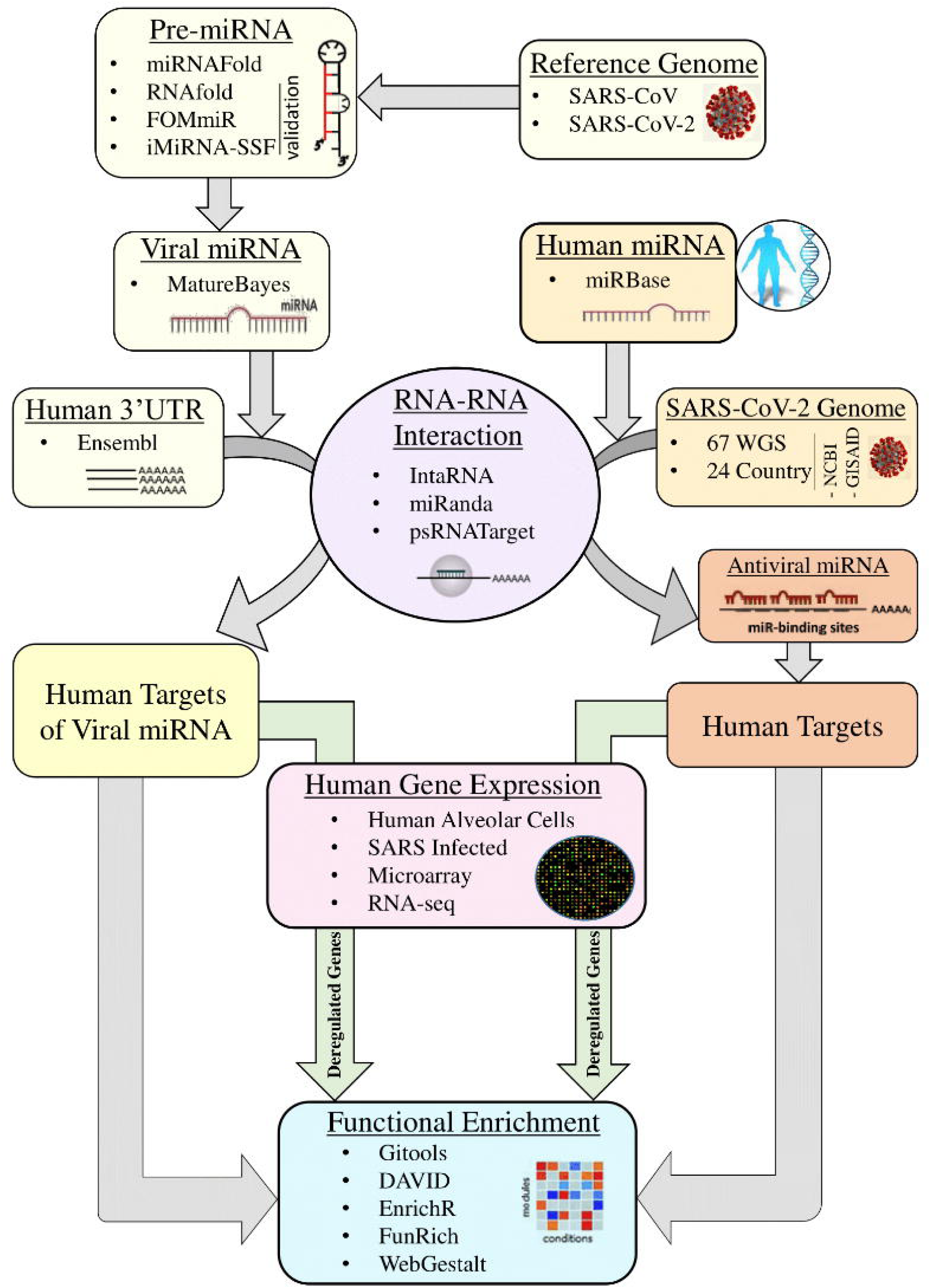
Overall workflow of the whole study.

## 2. Materials and Methods

### 2.1 Obtaining SARS-CoV and SARS-CoV2 Genome sequences

The reference genome of SARS-CoV (RefSeq Accession no. NC_004718.3) and SARS-CoV-2 (RefSeq Accession no. NC_045512.2) was fetched from NCBI RefSeq database (NCBI-RefSeq, 2020). Total 67 whole-genome sequences of SARS-CoV-2 isolates covering 24 different countries (Supplementary file 1) were retrieved from NCBI Virus (NCBI-Virus, 2020) and GISAID (GISAID, 2020).

### 2.2 Obtaining human 3’UTR and mature miRNA sequences

Human miRNAs were accessed from microRNA database miRBase (Kozomara et al., 2018) and 3’UTR sequences of human protein coding genes were obtained from Ensembl-Biomart (Hunt et al., 2018) (release 99).

### 2.3 Prediction of viral pre-miRNA and validation

We used miRNAFold (Tav et al., 2016) for *de novo* prediction of all possible precursor-miRNAs from the obtained reference sequences of SARS-CoV and SARS-CoV-2 with sliding window size of 150 and minimum hairpin size as 0. The results were validated using three different tools. First RNAfold (Gruber et al., 2008) was used with minimum free energy (MFE) and partition function fold algorithm to find stable secondary structures. Second, a fixed-order Markov model-based algorithm namely FOMmiR (Shen et al., 2012) was used and finally a SVM-based tool iMiRNA-SSF (Chen et al., 2016) was used that calculates minimum free energy (MFE), p-value of randomization test (P-value) and the local triplet sequence-structure features. The common predictions from these three tools were utilized for further analysis.

### 2.4 Prediction of mature miRNA

A Naive Bays classifier algorithm implemented in tool MatureBayes (Gkirtzou et al., 2010) was used to identify mature miRNA candidates within the miRNA precursor sequences.

### 2.5 RNA-RNA interaction analysis

Three different tools were used to analyze RNA-RNA interactions for host miRNA-viral genome and viral miRNA-host 3’UTR of coding sequences. IntaRNA 2.0 (Mann et al., 2017) was used considering sites with parameters --mode=H --model=X, --outMode=C, ΔΔG □≤ − 10□kcal/mol, with seed 2–8 allowing G:U base pairs. microRNA.org (Betel et al., 2008) was used with a score cutoff ≥□140, energy cutoff ≤ □ − 20kcal/mol, gap opening □=□ − 9.0 and gap extension□ = □− 4.0; psRNATarget (Dai and Zhao, 2011) was also used to determine RNA-RNA interactions. Finally, the common predictions from these three tools were considered for downstream analysis.

### 2.6 Target genes functional enrichment analysis

#### 2.6.1 Enrichment analysis in Gitools

The functional annotation of target genes is based on Gene Ontology (GO) (Ashburner et al., 2000) as extracted from EnsEMBL (Hubbard et al., 2007) and KEGG pathway database (Kanehisa and Goto, 2000). Accordingly, all genes are classified into the ontology categories’ biological process (GOBP) and pathways when possible. We have taken only the GO/pathway categories that have at least 10 genes annotated. We used Gitools for enrichment analysis and heatmap generation (Perez-Llamas and Lopez-Bigas, 2011). Resulting p-values were adjusted for multiple testing using the Benjamin and Hochberg’s method of False Discovery Rate (FDR) (Benjamini and Hochberg, 1995).

#### 2.6.2 Enrichment analysis using web-based tools

The host miRNA targeting SARS-CoV and SARS-CoV-2 were used for functional over-representation analysis to visualize and predict the roles of these miRNA in human diseases and find enriched pathways. Besides Gitools, functional enrichment analyses target human genes were conducted using; EnrichR (Kuleshov et al., 2016); DAVID 6.8 (Huang et al., 2009; Sherman and Lempicki, 2009); WebGestalt 2019 (Liao et al., 2019) and FunRich 3.1.3 (Pathan et al., 2017). The targeted genes are analyzed to determine their role in viral pathogenesis, infectivity, and immune evasion.

### 2.7 Microarray expression data analysis

Microarray data for change in gene expression induced by SARS-CoV on 2B4 cells infected with SARS-CoV or remained uninfected for 12, 24, and 48hrs obtained from Gene Expression Omnibus (GEO), ID GSE17400 (https://www.ncbi.nlm.nih.gov/geo) (Barrett et al., 2012). Raw Affymatrix CEL files were background corrected, normalized using Bioconductor package “affy” (version 1.28.1) using ’rma’ algorithm. Quality of microarray experiment (data not shown) was verified by Bioconductor package “arrayQualityMetrics” (Kauffmann et al., 2009) (version 3.2.4 under Bioconductor version 3.10; R version 3.6.0). To determine genes that are differentially expressed (DE) between two experimental conditions, Bioconductor package Limma (Smyth, 2005) was utilized to generate contrast matrices and fit the corresponding linear model. Probe annotations to genes were done using the Ensembl gene model (Ensembl version 99) as extracted from Biomart (Flicek et al., 2007) and using in-house python script. When more than one probes were annotated to the same gene, the highest absolute expression value was considered (maximizing). To consider a gene is differentially expressed, multiple tests corrected, FDR (Benjamini and Hochberg, 1995) p-value ≤ 0.05 was used as a cut-off.

### 2.8 RNA-seq expression data analysis

RNA-seq raw read-count data on SARS-CoV-2 mediated expression changes in primary human lung epithelium (NHBE) and transformed lung alveolar (A549) cells were obtained from the GEO database (GSE147507) (Barrett et al., 2012). For differential expression (DE) analysis we used Bioconductor package DESeq2 (version 1.38.0) (Anders and Huber, 2010) with R version 3.6.0 (Team, 1999) with a model based on the negative binomial distribution. To avoid false positive, we considered only those transcripts where at least 10 reads are annotated and a p-value of 0.01.

### 2.9 MicroRNA Clustering

The hierarchal clustering of human miRNAs that could target SARS-CoV-2 genomes (binary mode) obtained from various countries was done using Manhattan distance and complete linkage analysis with the Genesis tool (Sturn et al., 2002). Human death number (per million population) due to SARS-CoV-2 infection was obtained on 2^nd^ April 2020 from ‘worldometer’ website (Worldometer, 2020).

### 2.10 Overlap Analysis

Two or three-way overlap analysis was done using online venn-diagram program Venny 2.1.0 (Oliveros, 2018). In case of multiple pairwise overlaps and correlation analysis, as well as heatmap generation, were done using Gitools (Perez-Llamas and Lopez-Bigas, 2011).

### 2.11 Data Visualization

We have visualized human miRNA that binds to the virus genome in web-genome browsers NCBI genome data viewer (NCBI’s-genome-browser, 2020).

## 3. Results

### 3.1 Several human miRNAs are found to target SARS-CoV and SARS-CoV-2

It is possible that during viral infections, host-encoded miRNAs can modulate viral infections as a means of host immune response (Girardi et al., 2018). To identify possible host miRNAs that can get induced during the SARS-CoV (R) and SARS-CoV-2 (R) infections, we have utilized a bioinformatics approach. From our rigorous analysis pipeline which covers three different well-established algorithms (IntaRNA, miRanda, and psRNATargets) to predict RNA-RNA interactions, we have identified 122 and 106 host antiviral miRNAs against SARS-CoV (R) and SARS-CoV-2 (R), respectively (Figure 2A, 2B) (Supplementary file 2). Amongst these, 27 miRNAs were found to be targeting both viruses (Figure 2A). Whilst comparing these miRNAs with the antiviral miRNAs from VIRmiRNA (Qureshi et al., 2014), we have found 4 (hsa-miR-654-5p, hsa-miR-198, hsa-miR-622, hsa-miR-323a-5p) and 3 (hsa-miR-17-5p, hsa-miR-20b-5p, hsa-miR-323a-5p) host miRNAs against SARS-CoV (R) and SARS-CoV-2 (R), respectively, to have experimental evidence of having antiviral roles during infections (Figure 2A, 2B, 2C).

**Figure 2:**
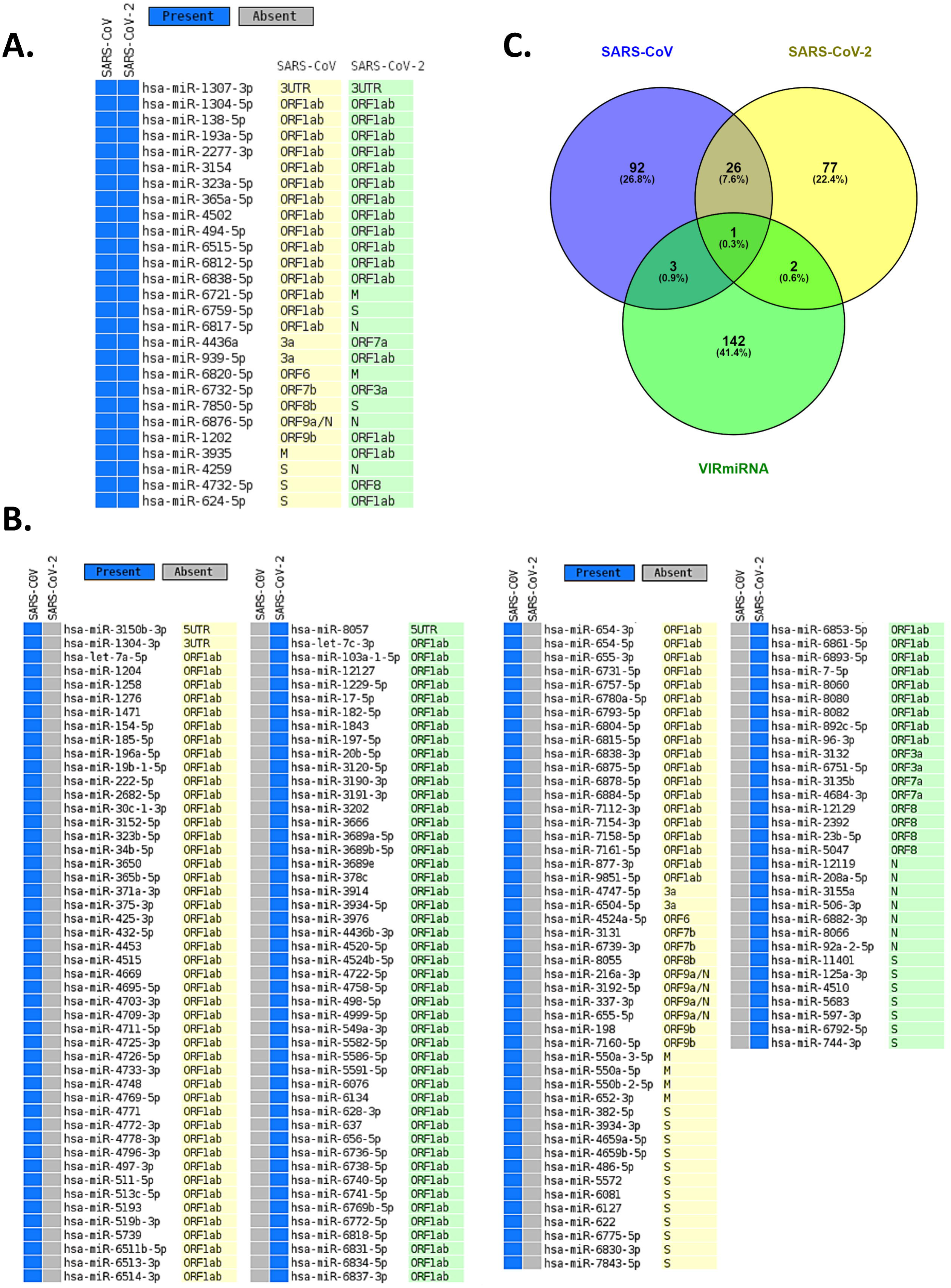
Virally induced host miRNAs targeting SARS-CoV and SARS-CoV-2. **A**. Common host miRNAs and their target genes in SARS-CoV and SARS-CoV-2, **B**. Host miRNAs and their target genes which uniquely target either SARS-CoV or SARS-CoV-2, **C**. Venn diagram showing the common and unique host miRNAs targeting SARS-CoV and SARS-CoV-2, and host miRNAs that have experimental evidences as antiviral miRNA.

Moreover, we compared the miRNAs targeting the two reference genomes of SARS-CoV (R) and SARS-CoV-2 (R) and found most of the host miRNAs can target the ORF1ab region, followed by the S region as the second-most targeted (Figure 3A, 3B). Also, the M, N, ORF3a, ORF7a, ORF8 (ORF8a, ORF8b for SARS-CoV), 5’ UTR and 3’ UTR regions of both viruses were targeted by host miRNAs. The significant variance was observed in the targeting positions of the host miRNAs between these two viruses (Figure 3A, 3B).

**Figure 3:**
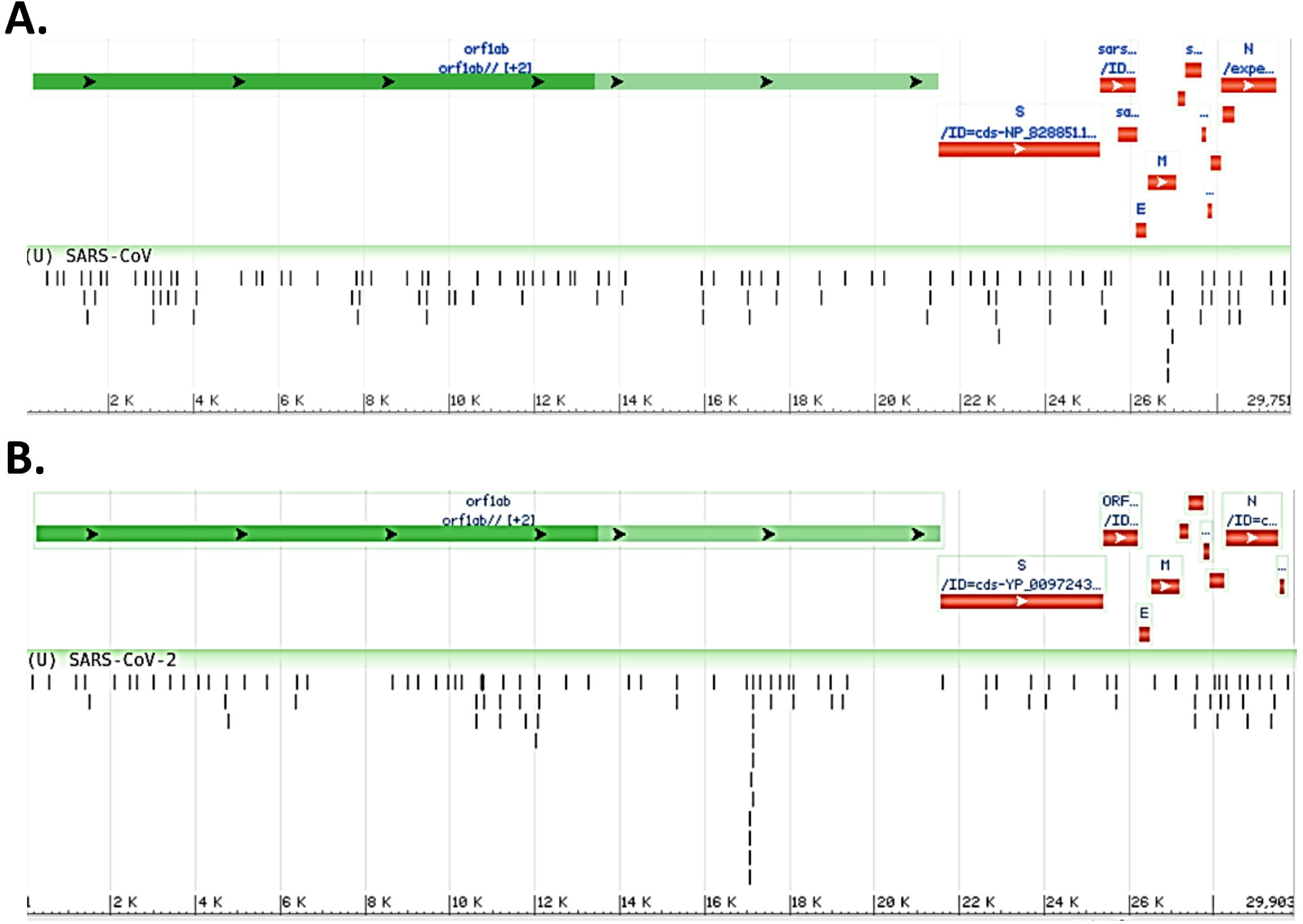
Genome browser view of host miRNAs targeting the regions of **A**. SARS-CoV (Reference) and **B**. SARS-CoV-2 (Reference) genomes.

Since RNA virus mutates fast, it is conceivable that mutations in crucial genomic locations would lead to differences in host miRNA binding patterns. Therefore, the ability of the host miRNAs targeting genomes of 67 SARS-CoV-2 isolates covering 24 different countries was also performed. Although, as expected, most of the identified host miRNAs’ binding profiles across these isolates remained somewhat similar to that of SARS-CoV-2 reference sequence; interestingly, we have identified 24 host miRNAs that bind differentially across the isolates (Figure 4A) which might have occurred due to the genomic variations between these isolates. Complete linkage agglomerative hierarchal cluster (HCL) analysis with Manhattan distance of these miRNAs (binary mode, bind or not bind) revealed two major clusters with a side cluster for one South Korean and two Singaporean isolates (Figure 4B). As miRNA is crucial in both host defense and viral pathogenesis, to understand the significance of this cluster, we have also compared the host miRNA clusters with the death rate (normalized by per million population) from different countries. Surprisingly, relatively higher deaths are found to be more prominent in the European major clusters (right side cluster) compared to the other major cluster (left side), and also found much lower deaths in side clusters (Figure 4B).

**Figure 4:**
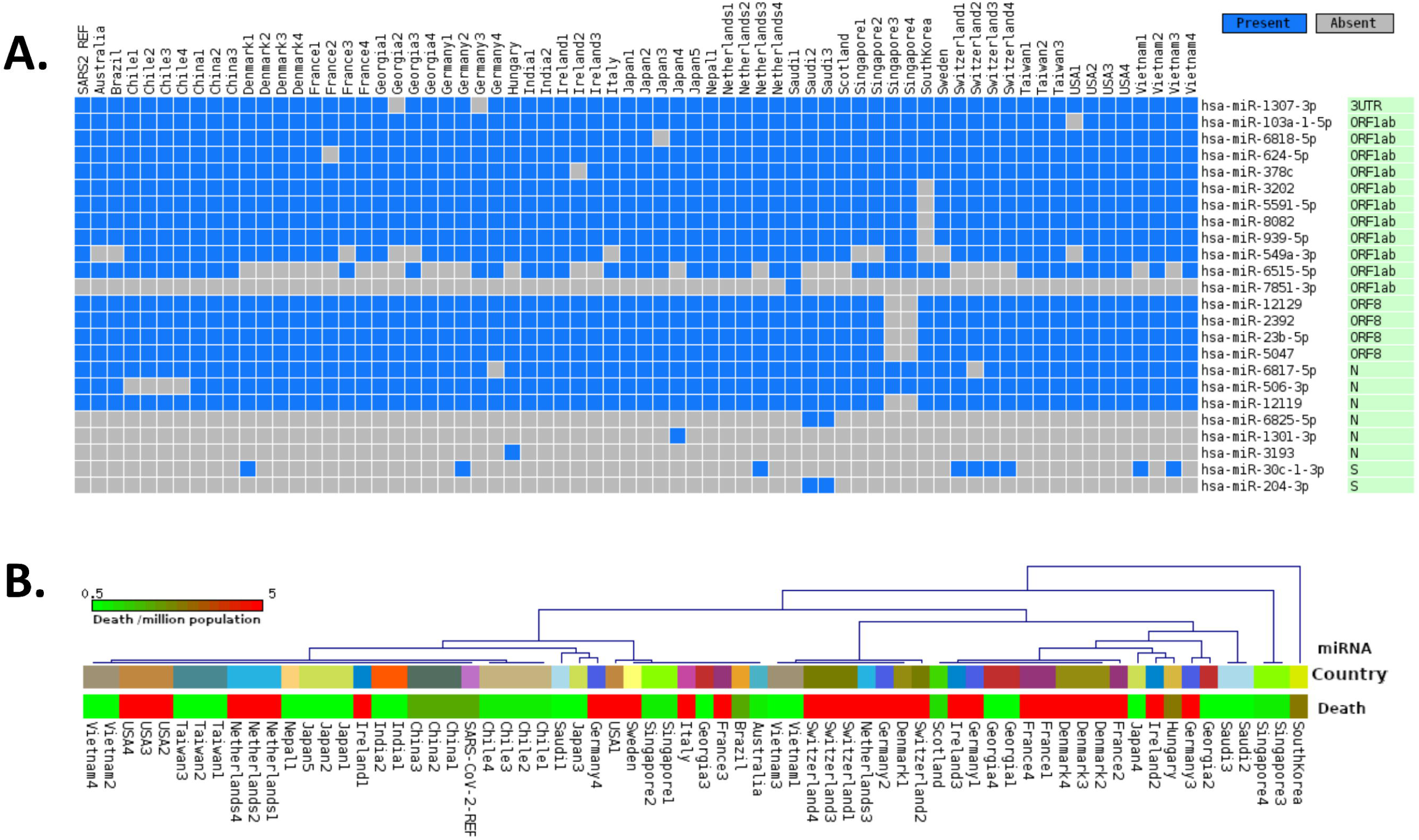
Differences of host miRNA binding profiles, **A**. representing only uncommon miRNAs binding pattern in 67 different SARS-CoV-2 genomes from 24 different countries, and **B**. Hierarchal clustering of all miRNAs binding in 67 genomes (upper panel, same country with same color code) and association of country specific death rates (in color coded scale) in per million population (lower panel).

### 3.2 Host miRNAs targeting SARS-CoV and SARS-CoV-2 play crucial roles in neutralizing the virus

Though the primary action elicited by host miRNAs is to silence the viral RNA, they might also modulate some host factors which provide an edge to the viral pathogenesis. To find out if these particular pathways are also targeted by the host miRNAs induced by SARS-CoV and SARS-CoV-2 infections, we have performed miRNA pathway enrichment analysis. We have found out several such pathways those might be deregulated by the host miRNAs to suppress the entry of the virus, to prevent the spread of the virions, and in minimizing the systemic symptoms resulting from the infection (Figure 5A).

**Figure 5:**
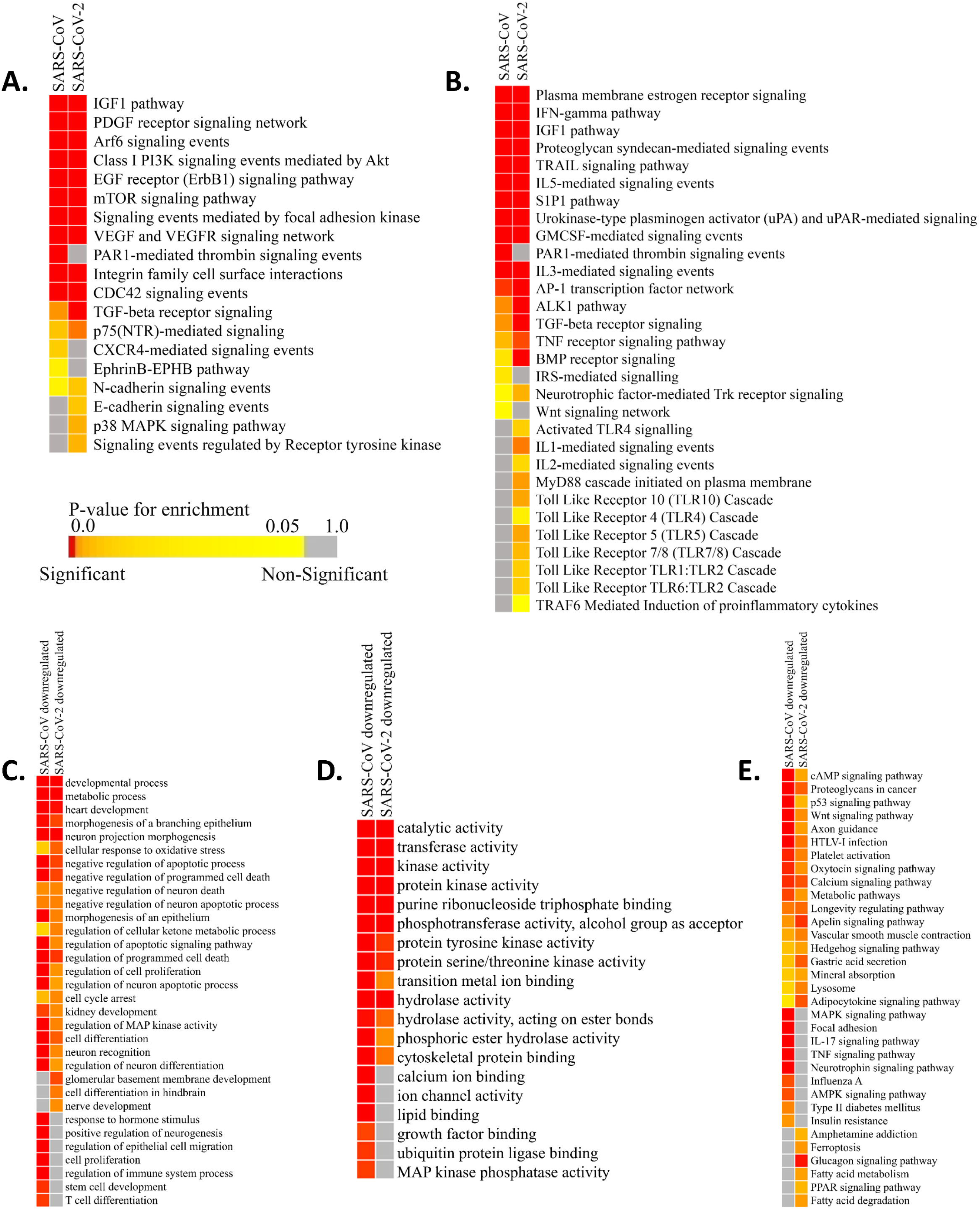
Enrichment analysis and comparison between host miRNA targets induced by SARS-CoV and SARS-CoV-2 infections. **A**. Heatmap representation of enriched pathways involved in host defense obtained using Funrich software, **B**. Enriched pathways which might act as proviral mechanisms obtained using Funrich software. Enrichment of downregulated host miRNA target genes in SARS-CoV and SARS-CoV-2 using gitools **C**. GO Biological Process module, **D**. GO Molecular Function module, **E**. KEGG pathway modules. Significance of enrichment in terms of adjusted p-value (< 0.05) is represented in color coded P-value scale for all heatmaps. Color towards red indicates higher significance and color towards yellow indicates less significance, while grey means non-significant.

Host miRNAs might have a probable role in blocking the entry of the virus, as they are found to be targeting the pathways needed for viral entry-PDGF receptor-like signaling (Soroceanu et al., 2008), Arf-6 signaling (García-Expósito et al., 2011), PI3K-Akt signaling (Diehl and Schaal, 2013), EGFR signaling (Zheng et al., 2014), signaling events mediated by focal adhesion kinase (Elbahesh et al., 2014), CDC42 signaling (Swaine and Dittmar, 2015), EphrinB-EPHB pathway (Wang et al., 2019), Cadherin signaling (Mateo et al., 2015), RTK signaling (Haqshenas and Doerig, 2019), etc (Figure 5A).

They can also block some machinery like-p38 MAPK signaling (Hirasawa et al., 2003), FAK signaling (Elbahesh et al., 2014), PI3K-Akt signaling (Diehl and Schaal, 2013), etc. which can be hijacked by viruses for their efficient replication, pre-mRNA processing and translation (Figure 5A). These host miRNAs might also try to reduce some host induced inflammatory responses to prevent acute lung damage by targeting IGF1 signaling (Li et al., 2019), VEGF signaling (Alkharsah, 2018), PAR1 signaling (Heuberger and Schuepbach, 2019), integrin signaling (Teoh et al., 2015), TGF-beta signaling (Denney et al., 2018), TRAIL signaling (Cummins and Badley, 2009), etc (Figure 5A). Some signaling pathways like-CXCR4 signaling (Arnolds and Spencer, 2014), TGF-beta signaling (Denney et al., 2018), mTOR signaling (Le Sage et al., 2016), PI3K-Akt signaling (Diehl and Schaal, 2013), etc. can facilitate viral survival in infected cells by inhibiting apoptosis, autophagy, early immune responses, etc. Host miRNAs may function to downregulate these to invoke a proper immune response against the viruses (Figure 5A).

### 3.3 Infection induced host miRNAs can function as a proviral factor by inhibiting host immune surveillance pathways

Host miRNAs can be like a double-edged sword as sometimes it can facilitate the viral immune evasion by targeting some important host immune responses (Bruscella et al., 2017). Our host miRNA enrichment analysis showed several significant pathways like-IFN-gamma signaling (Kang et al., 2018), TGF-beta signaling (Mogensen and Paludan, 2001), Interleukin signaling (Kimura et al., 2013), IGF1 signaling (Li et al., 2019), TRAIL signaling (Cummins and Badley, 2009), etc. which are involved in important proinflammatory cytokine signaling during viral infections (Figure 5B). Interestingly, we have found out that host miRNAs induced during SARS-CoV-2 infection may particularly downregulate different Toll-Like Receptors (TLRs) (Kimura et al., 2013) signaling which are considered as the primary stimulatory molecules for producing host antiviral responses (i.e. production of interferons and other inflammatory cytokines) (Figure 5B). Also, other receptor signaling involved in antiviral responses like-uPA-UPAR signaling (Alfano et al., 2003), TRAF6 signaling (Konno et al., 2009), S1P1 signaling (Oldstone et al., 2013), Estrogen receptor signaling (Kovats, 2015), Protease-activated Receptor (PAR) signaling (Antoniak et al., 2013), Bone morphogenetic protein (BMP) signaling (Eddowes et al., 2019), etc. can also be deregulated by the host miRNAs, leading to the host’s immune suppression (Figure 5B).

### 3.4 Host miRNAs’ targeted downregulated pathways are related to the comorbidities of COVID-19

SARS-CoV-2 infected patients with comorbidities (i.e. cardiovascular diseases, diabetes, renal problems) are found to be more susceptible to COVID-19. To find out whether virally induced host miRNAs are playing role in these, we have performed enrichment analysis using the downregulated targets genes of the host miRNAs using the expression data obtained from GEO dataset (GSE17400 for SARS-CoV and GSE147507 for SARS-CoV-2). These revealed that the downregulated targets of host miRNAs are involved in functions and pathways like-heart development, kidney development, several neuronal processes, metabolic process, regulation of cellular ketone metabolism, insulin resistance, glucagon signaling pathway, fatty acid metabolism, PPAR signaling, etc (Figure 5C, 5D, 5E). Aberrant regulation of these processes can overcomplicate the disease conditions of patients having existing disorders.

### 3.5 Viral miRNAs encoded by SARS-CoV and SARS-CoV-2 can target several host genes

Many human viruses were found to produce miRNAs to assist in their overall pathogenesis by modulating host factors (Bruscella et al., 2017). Previous study on SARS-CoV also suggests that viral small non-coding RNAs can help its efficient pathogenesis (Morales et al., 2017). Our bioinformatics approach suggests that SARS-CoV and SARS-CoV-2 can also encode some viral miRNAs. miRNAfold tool (Tav et al., 2016) yielded 529 and 519 putative pre-miRNAs from the genome of SARS-CoV and SARS-CoV-2, respectively. RNAfold tool (Gruber et al., 2008) predicted 303 and 308 of these precursors of SARS-CoV and SARS-CoV-2, respectively are highly stable for forming hairpin structures which is a prerequisite of mature miRNA formation. Using FomMiR (Shen et al., 2012) and IMiRNA-SSF (Chen et al., 2016), we then predicted which of these highly stable precursors can truly produce mature miRNAs. We have found 63 and 85 such precursors respectively for SARS-CoV and SARS-CoV-2. Using Maturebayes tool, from these precursors, we have identified 126 and 170 mature miRNAs from SARS-CoV and SARS-CoV-2, respectively (Supplementary file 3). We have predicted the human target genes by utilizing three different target prediction tools and to reduce false positive, we have taken only the common set. This returned 5292 and 6369 human target genes for SARS-CoV and SARS-CoV-2, respectively (Supplementary file 4). Out of these, 2992 genes are found to be common in both, while 2300 and 3377 genes were found to be unique targets of SARS-CoV and SARS-CoV, respectively. An apparent difference of the coding regions of miRNAs between SARS-CoV and SARS-CoV-2 was observed (Figure 6A, 6B).

**Figure 6:**
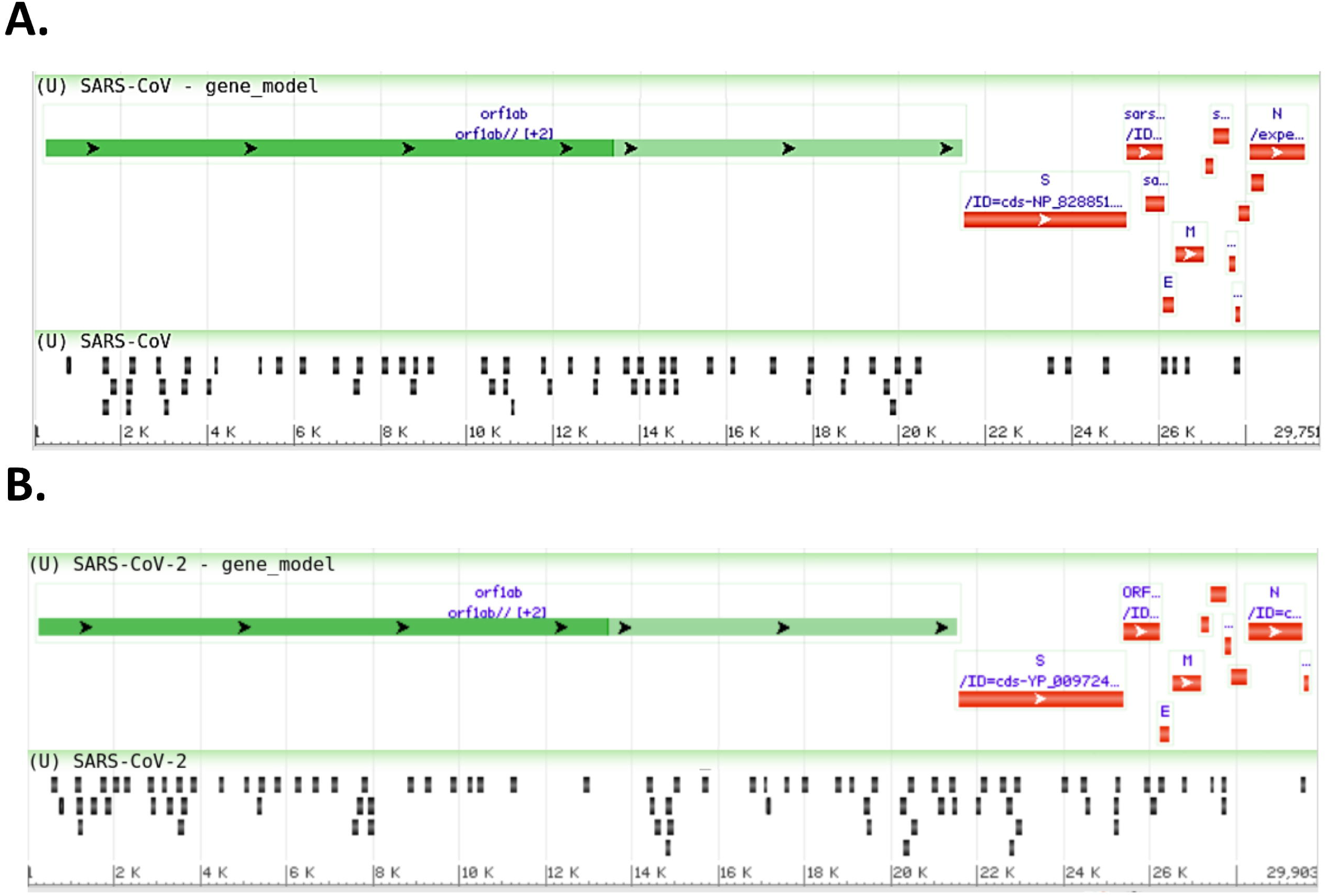
Genome browser view of viral miRNAs transcribed from the regions of **A**. SARS-CoV (Reference) **and B**. SARS-CoV-2 (Reference) genomes.

### 3.6 SARS-CoV and SARS-CoV-2 can evade host’s immune surveillance pathway by utilizing its miRNAs

Many viruses use their miRNAs to suppress or escape host’s immune responses (Mishra et al., 2020). To identify which pathways are associated with SARS-CoV and SARS-CoV infection, we have performed the gene ontology (GO) and pathway functional enrichment of the targeted genes using different tools. This reveals a myriad of significant functions and pathways involved in host immune responses, like-Wnt signaling (Ljungberg et al., 2019), MAPK signaling (Kimura et al., 2013), T cell-mediated immunity (Channappanavar et al., 2014), autophagy (Yordy and Iwasaki, 2011), FGF receptor binding (van Asten et al., 2018), TGF-beta signaling (Denney et al., 2018), VEGF signaling (Alkharsah, 2018), ErbB signaling (Zheng et al., 2014), mTOR signaling (Le Sage et al., 2016), TNF-alpha signaling (Kimura et al., 2013), etc are particularly targeted by SARS-CoV-2 (Figure 7A-7E).

**Figure 7:**
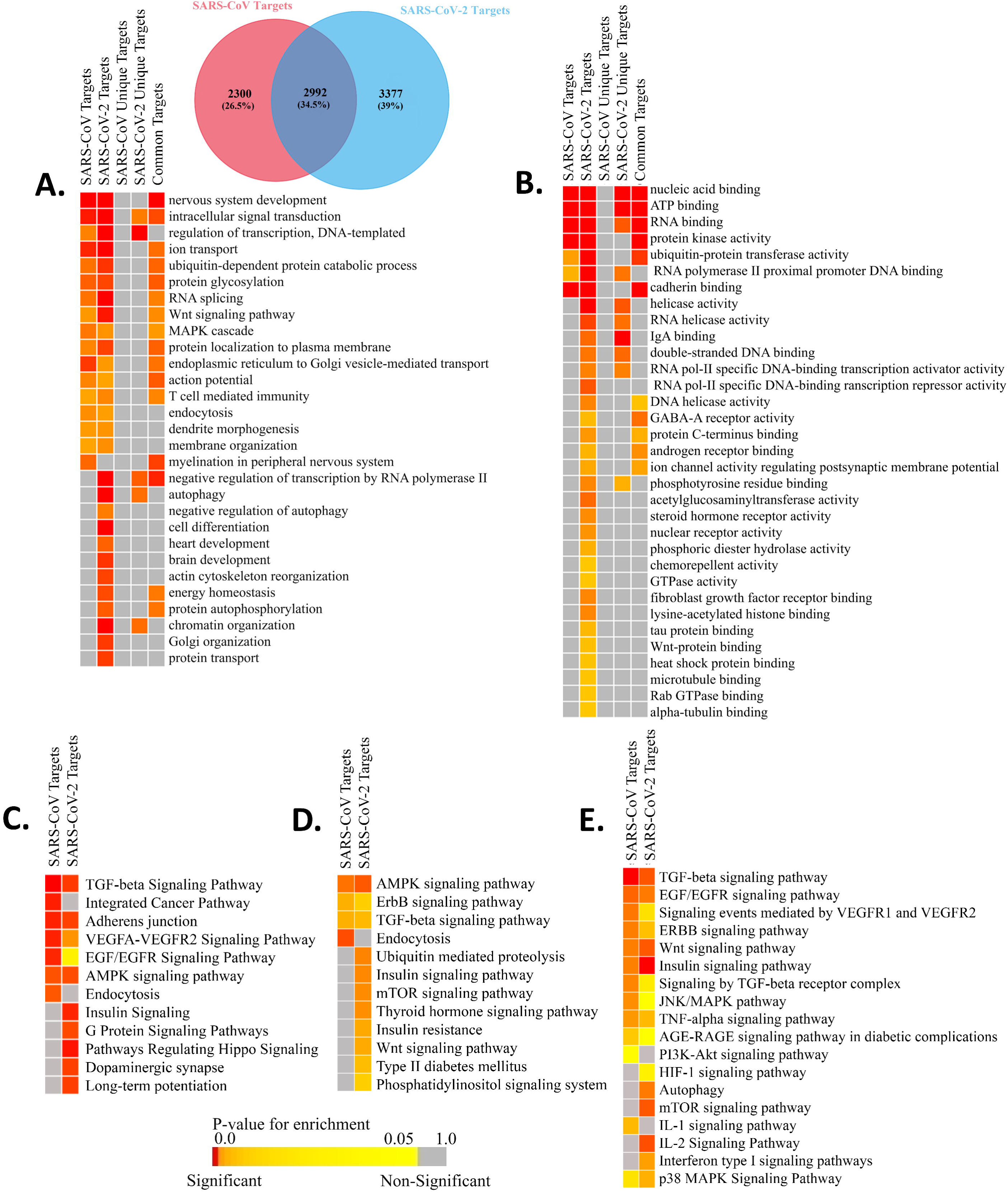
Enrichment analysis and comparison between the SARS-CoV and SARS-CoV-2 encoded viral miRNAs’ target human genes. Functional enrichment using gitools-**A**. GO Biological Process module, **B**. GO Molecular Funtion module. Enriched pathways obtained from-**C**. Webgestalt (KEGG and Wikipathways) tool, **D**. DAVID (KEGG pathways) tool, **E**. EnrichR (KEGG, Wikipathways, BioPlanet pathways) tool. Color codes are as in Figure 5.

Functions and pathways like heart development, brain development, and insulin signaling pathway, etc. (Figure 7A-7E) were also enriched for SARS-CoV-2 only, which can be targeted by the viral miRNAs, making the patients with previous complications more susceptible to COVID-19 as well as can lead to several signs uniquely found in SARS-CoV-2 infected patients.

We have also identified the downregulated target genes by curating the GEO expression datasets (GSE17400 for SARS-CoV and GSE147507 for SARS-CoV-2) and found 1890 and 35 downregulated target genes in SARS-CoV and SARS-CoV-2, respectively (Supplementary file 5). These downregulated target genes are found to be involved in different immune signaling pathways as well as different organ-specific functions related pathways (Figure 8).

**Figure 8:**
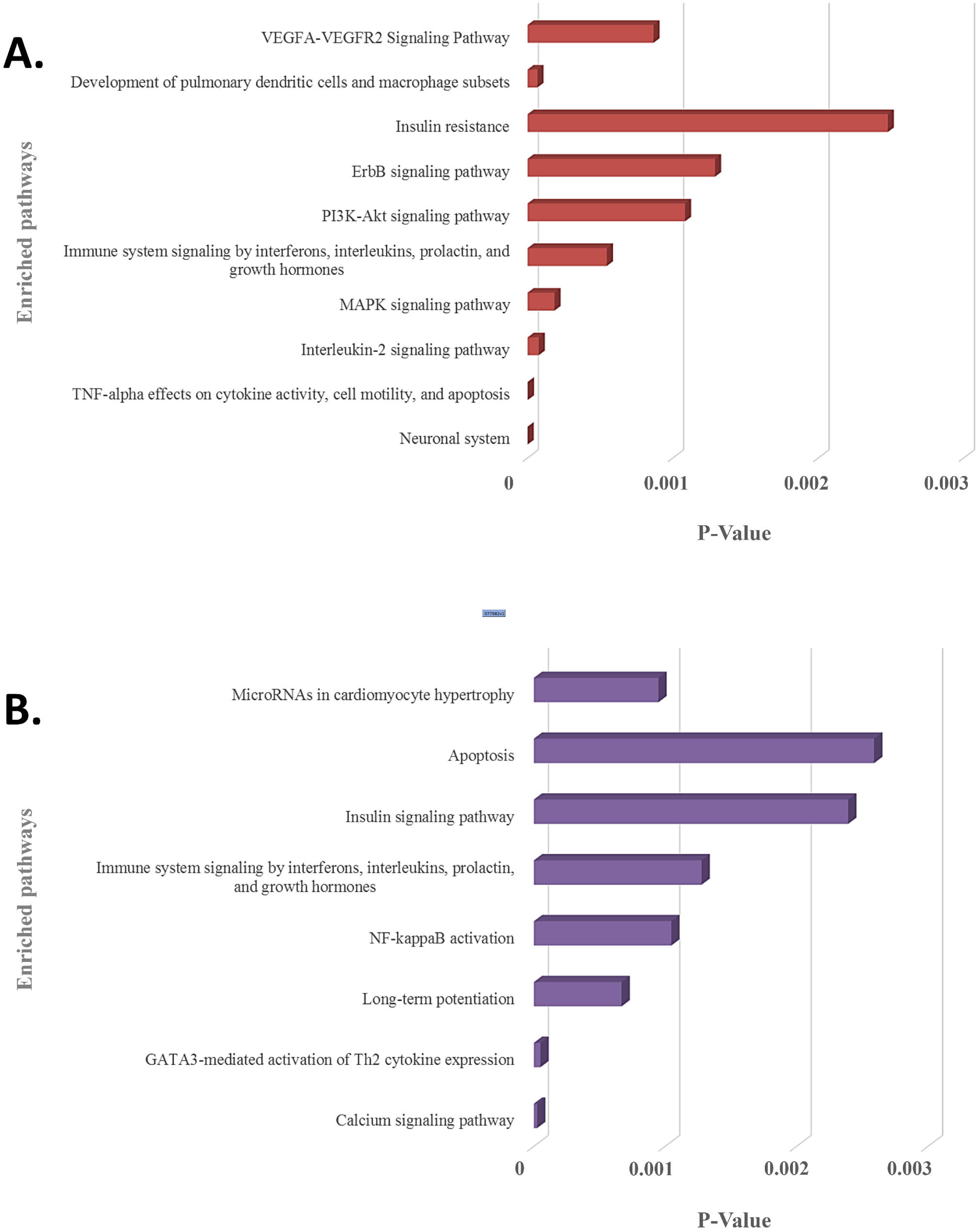
Enrichment analysis and comparison between the enriched pathways of **A**. SARS-CoV and **B**. SARS-CoV-2, encoded viral miRNAs’ downregulated target genes, obtained using EnrichR (KEGG, Wikipathways, BioPlanet pathways) tool. P-value scale is utilized for all in a bar graph. Higher the bar height, the more significant an enriched term is.

## 4. Discussion

Cellular miRNAs play a crucial role during the viral infection to strengthen host immunity by targeting virus’s genes as well as targeting pathways that viruses utilize for their survival and immune evasion (Girardi et al., 2018). Viruses themselves can encode their miRNAs to target these immune signaling pathways (Bruscella et al., 2017). COVID-19 has become a serious public health issue recently, though the complete molecular mechanism of pathogenesis is not fully understood yet. In this context, we have carried out this whole study to investigate the miRNA mediated interactions between host and SARS-CoV-2 virus, which might shed some insights on the tug-of-war between host’s immune responses and virus’s circumvention strategies. Though the disease conditions caused by SARS-CoV and SARS-CoV-2 are more or less similar, still several unique features (i.e. long incubation, enhanced latency, asymptomatic infection, intense pain, severe lung damage, etc. (Ceccarelli et al., 2020)) of SARS-CoV-2 making it more challenging to manage compared to SARS-CoV. So, we also sought to find out if there are any existing differences between SARS-CoV and SARS-CoV-2 in the context of miRNA mediated regulation of host responses.

As host miRNAs are one of the key immune protection against viral infections, we have tried to find out which cellular miRNAs can target SARS-CoV and SARS-CoV-2 genes. Due to differences in the genome sequences between these two viruses, there was a significant difference between cellular miRNAs and their targeting viral genes. Likewise, some of the commonly found cellular miRNAs were showing differential binding preferences for these viral genes. (Figure 2A). Previous study by Mallick *et al*. showed that cellular miRNAs can boost up host’s immune response as well as they can assist in viral immune evasion mechanisms (Mallick et al., 2009). Another study by Morales *et al*. suggested that SARS-CoV can encode small non-coding RNAs which can play roles in inflammatory lung pathology (Morales et al., 2017). We also compared the induced host miRNAs’ profiles of 67 SARS-CoV-2 isolates from 24 different countries across the globe. From this analysis, we have identified several clusters and associated miRNAs, and our correlation study between these clusters with the death counts all over the world shed some light on the burning question and suggests why the Europeans are more prone to COVID-19 (Figure 3B).

We found several miRNAs with experimentally validated antiviral roles; among those, hsa-miR-323a-5p and hsa-miR-654-5p (predicted for SARS-CoV) were found to inhibit viral replication in H1N1 Influenza virus infection (Song et al., 2010), while hsa-miR-17-5p and hsa-miR-20b-5p (predicted for SARS-CoV-2) were found to be upregulated in H7N9 Influenza virus infection (Zhu et al., 2014).

Apart from the basic role of cellular miRNAs in eliminating the transcripts of viruses, they can also modulate some host pathways which supposedly can be utilized by the infecting virus to avoid host’s immune response. We also identified several such pathways involved in viral entry, replication, translation mechanisms, etc. which can be targeted by the cellular miRNAs induced by SARS-CoV and SARS-CoV-2 infection. Moreover, several immune response pathways like-TLR signaling, interleukin signaling, TRAF6 signaling, etc. were exclusively found to be targeted by SARS-CoV-2 induced host miRNAs (Figure 5B) and SARS-CoV-2 encoded miRNAs can target pathways like-autophagy, IFN-I signaling, wnt signaling, mTOR signaling, etc., but SARS-CoV encoded miRNAs’ targets were not found to be enriched in these pathways (Figure 7A-E). Target genes downregulated by SARS-CoV-2 miRNAs are found to be involved in Ca^2+^ signaling pathway which is considered important activators of many signaling pathways (Zhou et al., 2009) (Figure 8B). All of these suggest why SARS-CoV-2 infections might be fatal for those who are immunosuppressed (D’Antiga, 2020).

Interestingly, our findings have enlightened several poorly understood mechanisms behind many of the unique clinical and pathological features of SARS-CoV-2 which has made it significantly different from SARS-CoV. Our predicted both cellular miRNAs and viral encoded miRNAs, induced during SARS-CoV and SARS-CoV-2 infection, were found to target cytokine signaling pathways involved in immune responses leading to the improved viral pathogenesis. Also, we found that SARS-CoV-2 miRNAs can target different important organ specific cellular functions and pathways. We showed that SARS-CoV-2 encoded miRNAs can target insulin signaling pathway (Figure 7A, Supplementary figure 1) and aberration of this pathway might overcomplicate the whole disease condition for COVID-19 patients with existing diabetic problems (Shimizu et al., 1980; del Campo et al., 2012). Our data also suggests that the SARS-CoV-2 miRNAs can target heart development-related pathways (Figure 7A, Supplementary figure 1), which might lead to similar consequences like viral myocarditis (Dennert et al., 2008) making the disease more fatal for the patients with existing cardiovascular complications. These SARS-CoV-2 encoded miRNAs might also target genes associated with brain development (Figure 7A, Supplementary figure 1) which might provide clue about the neurological signs like-headaches, vomiting, and nausea. SARS-CoV-2 induced host miRNAs can also downregulate kidney development and regulation of cellular ketone metabolic processes etc. (Figure 5C) increasing kidney’s burden, (Kanikarla-Marie and Jain, 2016) which might be fatal for patients who have diabetes and kidney complications. HIF-1 signaling was also found to be targeted by SARS-CoV-2 miRNAs (Figure 7E, Supplementary figure 1). This pathway is found to be associated with many viral infections as HIF-1 plays an important role in cellular survival during hypoxic conditions (Santos and Andrade, 2017); COVID-19 patients suffer from the lack of oxygens due to breathing complications; so this pathway might play crucial roles to mitigate the condition, but viral miRNA mediated deregulation of this pathway might result in severe consequences.

Our findings can explain that the interplay of miRNAs of host and SARS-CoV-2 virus can promote viral pathogenesis by deregulating major antiviral immune signaling pathways, as well as abnormal regulations of several host pathways, might lead to increased complications in the infected patients. Our study that is conducted using machine learning and knowledgebase approaches, with further experiments, has the full potential to provide a more detailed understanding of the disease progression and based on these results, novel therapeutic interventions using RNA interference (RNAi) can be designed.

## Author’s contribution

ABMMKI conceived the project. ABMMKI and MAAKK designed the workflow. MAAKK, MRUS, MSI and MSM collected the data. All authors performed the analyses. MAAKK, MRUS, MSI and ABMMKI wrote the manuscript. All authors read and approved the final manuscript.

## Acknowledgments

We acknowledge Rafeed Rahman Turjya for valuable suggestions.

## Data Availability Statement

Publicly available data were utilized. Analyses generated data are deposited as supplementary files.

## Supplementary Figure Legend

**Supplementary figure 1:** SARS-CoV-2 miRNA targeted Genes involved in significant functions/pathways- **A**. Autophagy (Figure 7A, 7E), **B**. Heart development (Figure 7A), **C**. Brain development (Figure 7A), **D**. Insulin signaling pathway (Figure 7C, 7D, 7E), **E**. Interferon type 1 signaling pathway (Figure 7E), **F**. HIF-1 signaling pathway (Figure 7E).

## List of Supplementary files

**Supplementary file 1:** List of SARS-CoV-2 isolates used.

**Supplementary file 2:** List of Human miRNAs targeting SARS-COV (Reference) and SARS-CoV-2 (Reference) genes.

**Supplementary file 3:** List of predicted viral miRNAs of SARS-CoV (Reference) and SARS-CoV-2 (Reference).

**Supplementary file 4:** List of human target genes targeted by SARS-CoV (Reference) and SARS-CoV-2 (Reference) encoded miRNAs.

**Supplementary file 5:** Downregulated human target genes of SARS-CoV (Reference) and SARS-CoV-2 (Reference) encoded miRNAs.

