## Supplementary figures and images for "Epigenetic regulator miRNA pattern differences among SARS-CoV, SARS-CoV-2 and SARS-CoV-2 world-wide isolates delineated the mystery behind the epic pathogenicity and distinct clinical characteristics of pandemic COVID-19"

### Supplementary Figure 1

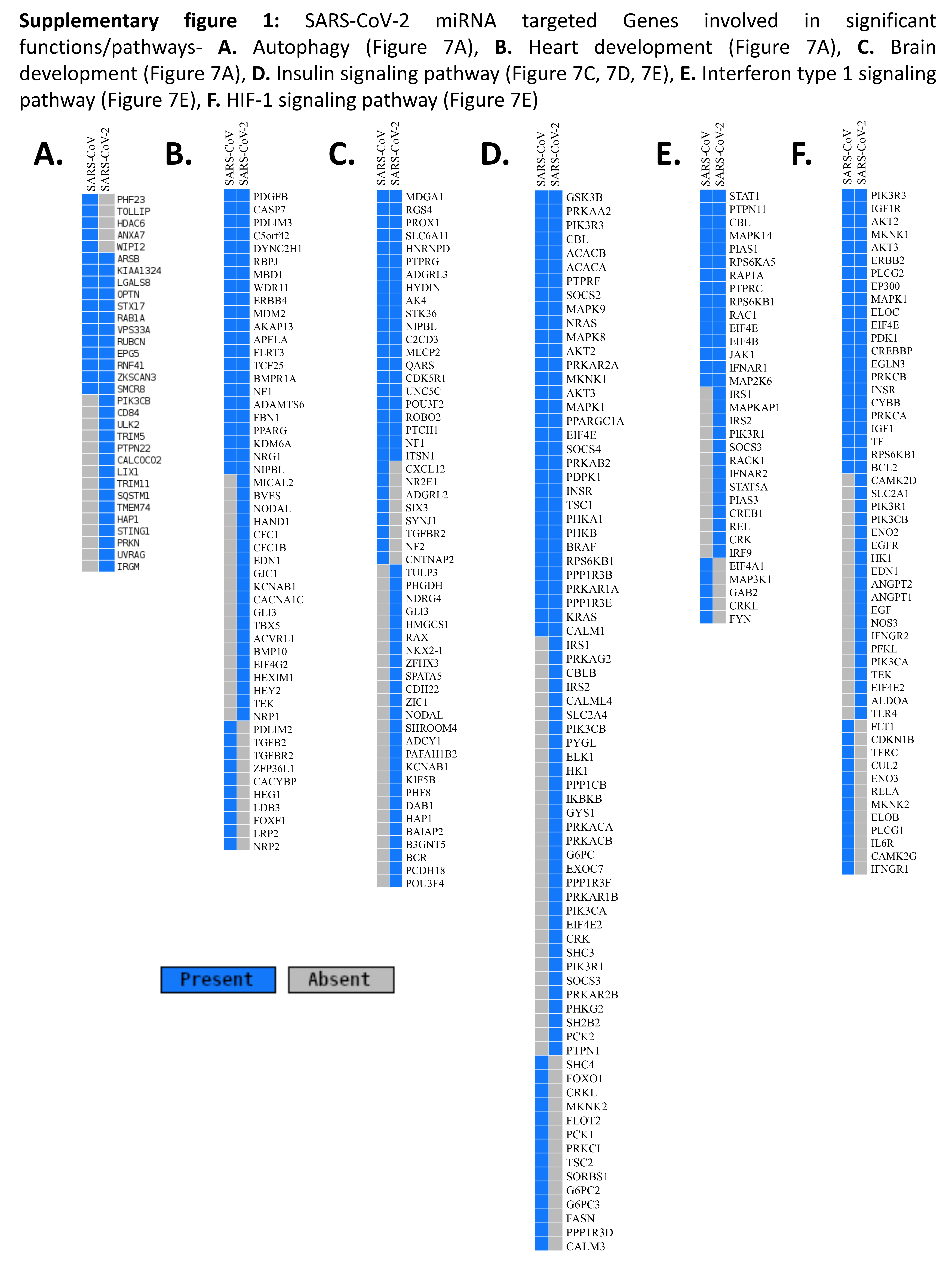
